# Initiation and regulation of vascular tissue identity in the *Arabidopsis* embryo

**DOI:** 10.1101/832501

**Authors:** Margot E. Smit, Cristina I. Llavata-Peris, Mark Roosjen, Henriette van Beijnum, Daria Novikova, Victor Levitsky, Daniel Slane, Gerd Jürgens, Victoria Mironova, Siobhan M. Brady, Dolf Weijers

## Abstract

Development of plant vascular tissues involves tissue specification, growth, pattern formation and cell type differentiation. While later steps are understood in some detail, it is still largely unknown how the tissue is initially specified. We have used the early Arabidopsis embryo as a simple model to study this process. Using a large collection of marker genes, we find that vascular identity is established in the 16-cell embryo. After a transient precursor state however, there is no persistent uniform tissue identity. Auxin is intimately connected to vascular tissue development. We find that while AUXIN RESPONSE FACTOR5/MONOPTEROS/ (ARF5/MP)-dependent auxin response is required, it is not sufficient for tissue establishment. We therefore used a large-scale enhanced Yeast One Hybrid assay to identify potential regulators of vascular identity. Network and functional analysis of suggest that vascular identity is under robust, complex control. We found that one candidate regulator, the G-class bZIP transcription factor GBF2, modulates vascular gene expression, along with its homolog GBF1. Furthermore, GBFs bind to MP and modulate its activity. Our work uncovers components of a gene regulatory network that controls the initiation of vascular tissue identity, one of which involves the interaction of MP and GBF2 proteins.

## Introduction

Vascular tissues play a central role in plant growth and development by providing plants with both transport capabilities and structural support. Steps in development of vascular tissues have been studied in detail, mainly in the Arabidopsis leaf (Donner et al., 2009; Gardiner et al., 2011; Krogan et al., 2012), shoot (Etchells et al., 2013; Hirakawa et al., 2010; Smetana et al., 2019; McConnell et al., 2001; Han et al., 2018) and root (Scheres et al., 1995; Miyashima et al., 2019; De Rybel et al., 2014). From this wealth of studies, a picture emerges where dedicated regulatory modules function to create a properly sized and patterned transport bundle. Several steps can be recognized in this process: specification of vascular tissue identity, cell proliferation to generate a bundle of cells, patterning into xylem, phloem and cambium cell types and finally differentiation into functional transport cells. The regulators and effectors of all but the first step have been dissected in some detail.

The rate of proliferation by periclinal cell divisions determines the width of a vascular bundle. Periclinal cell divisions in the vascular cells are controlled by several pathways: one directed by the xylem expressed TARGET OF MONOPTEROS 5 (TMO5) - LONESOME HIGHWAY (LHW) dimer (De Rybel et al., 2013, 2014; Ohashi-Ito et al., 2014, 2013), another regulated by the phloem-expressed PHLOEM EARLY DOFs (PEARs) (Miyashima et al., 2019), and a third depending on the activity of WUSCHEL-LIKE HOMEOBOX 4 and 14 in the cambium (WOX4/14) (Etchells et al., 2013; Fisher and Turner, 2007; Hirakawa et al., 2010). In concert with proliferation, cells in the vascular bundle develop a pattern of distinct sub-identities. Xylem development is associated with high auxin signaling, and further specification of proto- or metaxylem identity depends on a combination of cytokinin response and the activity of HD-ZIP III transcription factors (Baima et al., 2001; Bishopp et al., 2011; Carlsbecker et al., 2010; Mähönen et al., 2006; McConnell et al., 2001). Conversely, determination of phloem identity is associated with high cytokinin activity and the presence of, among others, ALTERED PHLOEM DEVELOPMENT (APL) (Bonke et al., 2003). Located between the phloem and xylem, the meristem-like (pro)cambium was shown to contribute to both the xylem and the phloem cell populations (Smetana et al., 2019). Finally, several vascular cell types undergo irreversible, terminal differentiation. The differentiation of xylem vessel elements can be triggered when a gene regulatory network under control of VASCULAR-RELATED NAC-DOMAIN6 (VND6) and VND7 is initiated (Kubo et al., 2005; Yamaguchi et al., 2010; McCarthy et al., 2009). While no differentiation-inducing factor has yet been found to trigger phloem-like differentiation (Blob et al., 2018), several factors necessary for phloem differentiation have been identified (Ruiz Sola et al., 2017; RodriguezVillalon et al., 2014; Wallner et al., 2017).

Most of the studied regulators of vascular development are expressed only or preferentially in vascular cells, which suggests the existence of a robust genetic identity. However, it has so far remained elusive how this vascular tissue identity is triggered or established. *De novo* vascular identity specification occurs repeatedly during the life cycle as new organs develop or when tissues are wounded (León et al., 2001; Melnyk et al., 2015; Yin et al., 2012). Specification of tissue identities involves the local accumulation of a signaling molecule (small molecule, peptide or protein) that will either promote or suppress the activation of a cell type-specific gene regulatory network. Such mechanisms have been described in nonhair versus hair cells in the root (Lee and Schiefelbein, 1999; Bernhardt et al., 2005), meristemoids versus stomatal-lineage ground cells in the stomatal lineage (Zhang et al., 2016; Yang et al., 2015; Lau et al., 2014) and xylem versus phloem cells in the vascular bundle (Smetana et al., 2019; Mähönen et al., 2000; Baima et al., 2001).

A signaling molecule that is strongly correlated with vascular development is auxin. Vascular tissue can be initiated by a source of auxin in cut stems (Sachs, 1969), and as a result, auxin maxima are often associated with vascular development (Brackmann et al., 2018, Miyashima et al., 2019; Scarpella et al., 2006; Wabnik et al., 2013).c Conversely, lack of the AUXIN RESPONSE FACTOR5/MONOPTEROS (MP) transcription factor causes impaired vascular development in the Arabidopsis embryo, seedling, leaf and stem (Berleth and Jürgens, 1993; Hamann et al., 1999; Hardtke and Berleth, 1998; Mayer et al., 1991). Indeed, c MP controls a variety of vascular-specific genes and pathways (De Rybel et al., 2013; Donner et al., 2009; Möller et al., 2017; Schlereth et al., 2010; Yoshida et al., 2019). However, all reported perturbations of auxin activity (synthesis, transport, response) that affect vascular development also affect a range of other processes (Bennett et al., 1996; Cheng et al., 2006; Marchant, 1999; van den Berg and ten Tusscher, 2017). It is therefore difficult to separate a role for auxin in the initiation of vascular identity from its many other functions, which warrants the use of an simple developmental model system for studying vascular tissue initiation in the absence of e.g. differentiation. The early embryo is an attractive model given that it lacks confounding wound response or extensive proliferation. Its simplicity and predictable division pattern allows detection of early developmental defects (Scheres et al., 1994) and available transcriptome resources (Palovaara et al., 2017; Belmonte et al., 2013; Schon and Nodine, 2017; Slane et al., 2014) enables in-depth investigations of vascular identity establishment.

Here we first use a suite of established and novel transcriptional reporters to track the stepwise establishment of vascular tissue identity in the embryo. We find that the identity initially established is unique to the embryo, transitioning to a mature and robust identity in the root. We show that auxin response is necessary but not sufficient to establish vascular identity. Via large-scale enhanced Yeast One Hybrid assays we identify common regulators of vascular genes and we find that one of these, the bZIP transcription factor GBF2, can interact with ARFs and modifies MP activity in the regulation of vascular-specific genes.

## Results

### Establishment of vascular tissue identity is a multi-step process

As cell type identity is often instructed at the level of gene activity, it can be inferred by gene expression markers that are uniquely present in said cell type. Here, we used a large and diverse set of established cell type markers to ask when vascular tissue specification occurs during embryogenesis, and to determine the ontogeny of the tissue. We selected *SHR* and *ATHB8* as early vascular markers in root and leaf development (Baima et al., 2001; Gardiner et al., 2011; Nakajima et al., 2001). *ZLL* shows vascular-specific expression in the root and embryo (Radoeva et al., 2016; Tucker et al., 2008; Haseloff, 1999). *WOL, PEAR1* and *DOF6* are all associated with cytokinin-responsive growth in the vascular bundle of the root (Mähönen et al., 2000; Miyashima et al., 2019). In contrast, expression of *TMO5, T5L1, TMO6, IQD15, SOK1* and *WRKY17* was shown to depend on MP (Schlereth et al., 2010; Möller et al., 2017).

Lineage tracing has suggested that the first vascular cells are specified in the early globular stage embryo (Scheres et al., 1994)(Figure 1B), when the embryo first contains three distinct cell layers. Indeed, we found almost all vascular reporters to be expressed at this stage, with the exception of *TMO6* and *T5L1* (Figure 1A). However, at this stage, most reporters were expressed in both the central cell layer, and in the surrounding ground tissue cells: *ATHB8*, *DOF6*, *PEAR1*, *WOL* and *ZLL* are expressed at apparently equal levels in both tissue types. In contrast, *IQD15* and *SOK1* show only low levels of ground tissue expression (Figure 1A). The only markers that were restricted to the innermost cell layer are *TMO5* and *SHR*, whose expression could not be detected in the ground tissue (Figure 1A). The broad expression of these vascular marker genes is only transient: within several cell division rounds, all vascular markers are restricted to the innermost cells, a pattern that is maintained in the postembryonic root (Figure 1A). This suggests that, rather than being immediately restricted to a small number of innermost cells, vascular identity starts as a broad trait that is limited to the inner cells over time. While all 12 marker genes eventually become restricted to the vascular cells, it appears that there are multiple trajectories to their vascular-specific expression pattern.

**Figure 1:**
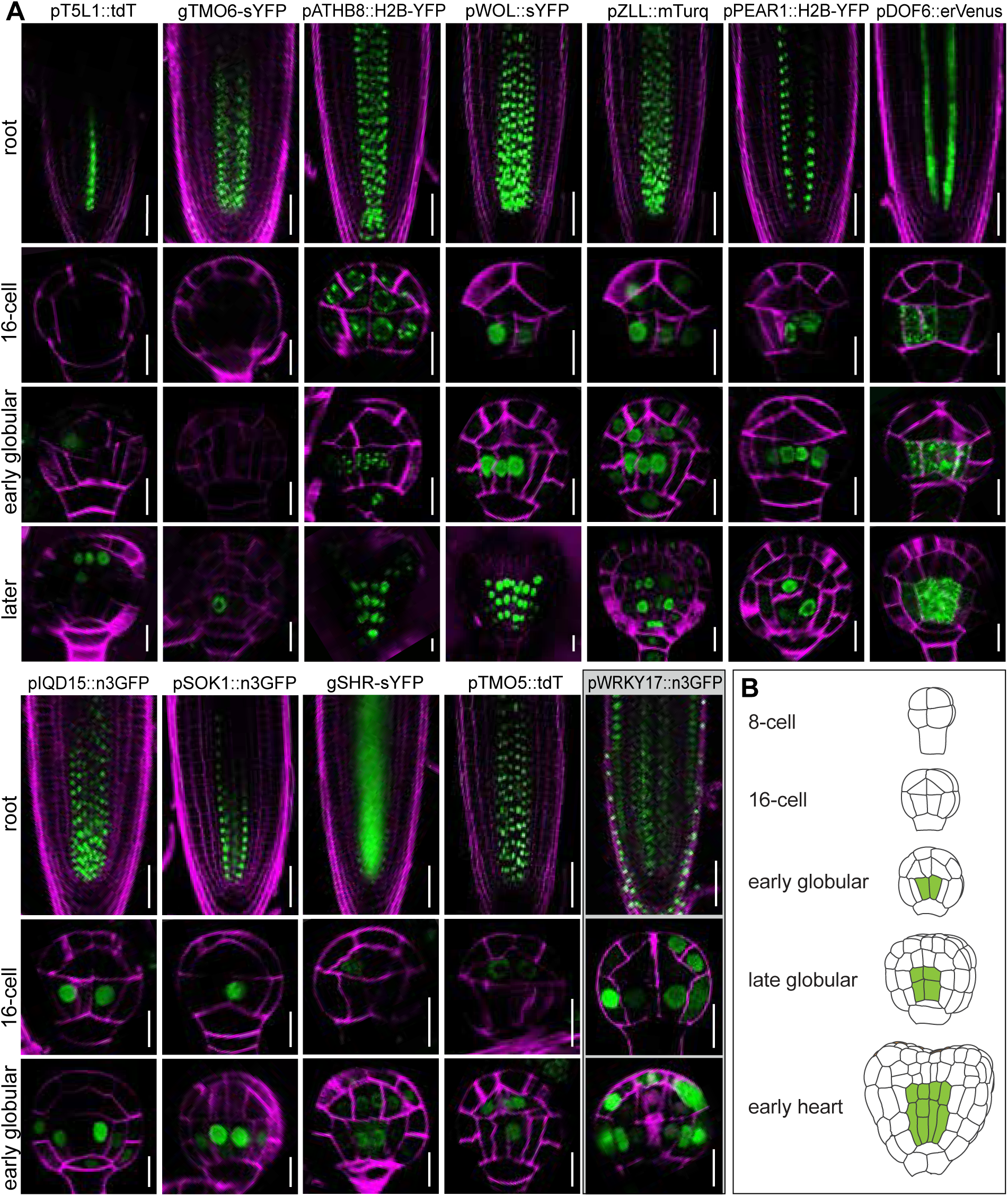
Expression patterns of previously described vascular reporters. (A) Expression patterns of previously published vascular reporters in root and early embryo. Insets shows 5 stages of embryogenesis. All reporters are transcriptional reporters except those for SHR and TMO6, which are translational fusions. Fluorescent protein signals are displayed in green, cell wall staining in magenta. Roots are stained with PI, embryos with Renaissance. Scale bars represent 50 µm (root) or 10 µm (embryo). (B) Overview of stages of early embryogenesis. Cells previously discussed as vascular are marked in green.

The route to a vascular-specific expression pattern starts in the 16-cell stage embryo rather than at early globular stage as previous reports have suggested. In previous work, inner lower tier cells at 16-cell stage were shown to resemble their vascular daughter cells at the globular stage using GO term enrichment in transcriptomes (Palovaara et al., 2017). Indeed, we find that many vascular marker genes start expression at 16-cell stage, where they are exclusively expressed in the inner cells: *DOF6*, *IQD15*, *PEAR1*, *SOK1*, *WOL* and *ZLL* (Figure 1A). Thus, as the 8-cell embryo divides to generate outer and inner cell layers in the 16-cell stage, inner cells activate vascular markers. Their ground tissue daughters initially retain the expression of some vascular markers, and switch these off later.

Many of the vascular marker genes were originally identified as targets of auxin signaling, often regulated by MP (Schlereth et al., 2010; Möller et al., 2017). As a result, there is a bias towards auxin-regulated genes among the well-studied vascular marker genes. We therefore searched for novel marker genes in an unbiased manner. Vascular-enriched genes were selected based on their expression in the early vascular cells, using a cell-type specific embryo transcriptome atlas (Palovaara et al., 2017) and additional publicly available vascular-specific transcriptome datasets (Figure 2A)(Brady et al., 2007; Belmonte et al., 2013; Kondo et al., 2015; Melnyk et al., 2018). Transcriptional reporter lines were constructed to test the expression pattern of 36 potential marker genes and eventually 5 qualified as markers of vascular identity during embryogenesis. Expression of the remaining 31 genes could either not be detected during embryogenesis, or was not limited to vascular cells in the embryo, whilst being specific to vascular tissue in the root (Figure S1). Of the 5 selected reporters, *GATA20* and *AP2B3* expression starts at 16-cell stage, and at the early globular stage, both are enriched in vascular cells (Figure 2B). *AP2B3* expression peaks in vascular cells but can also be detected in surrounding cell layers (Figure 2B), while *GATA20* expression shows vascular specificity (Figure 2B). In the root tip, *GATA20* has been shown to be expressed in the phloem (Lee et al., 2006) and we find that this expression is broader in the vascular cells close to the QC (Figure S1B). The other 3 selected reporters: *MEE45*, *MIR171B* and *MSS3*, are expressed at the dermatogen stage in all cells, but at lower levels in the vascular cells within the embryo, thus negatively marking vascular cells. Therefore, we will refer to these three as “inverse” markers of vascular identity in the embryo. This pattern is similar to that of *WRKY17*, a target of MP (Möller et al., 2017) (Figure 1A). However, while *WRKY17* is expressed broadly in the root meristem, *MEE45*, *MIR171B* and *MSS3* show tissue-specific expression in the root (Figure 2B).

**Figure 2.**
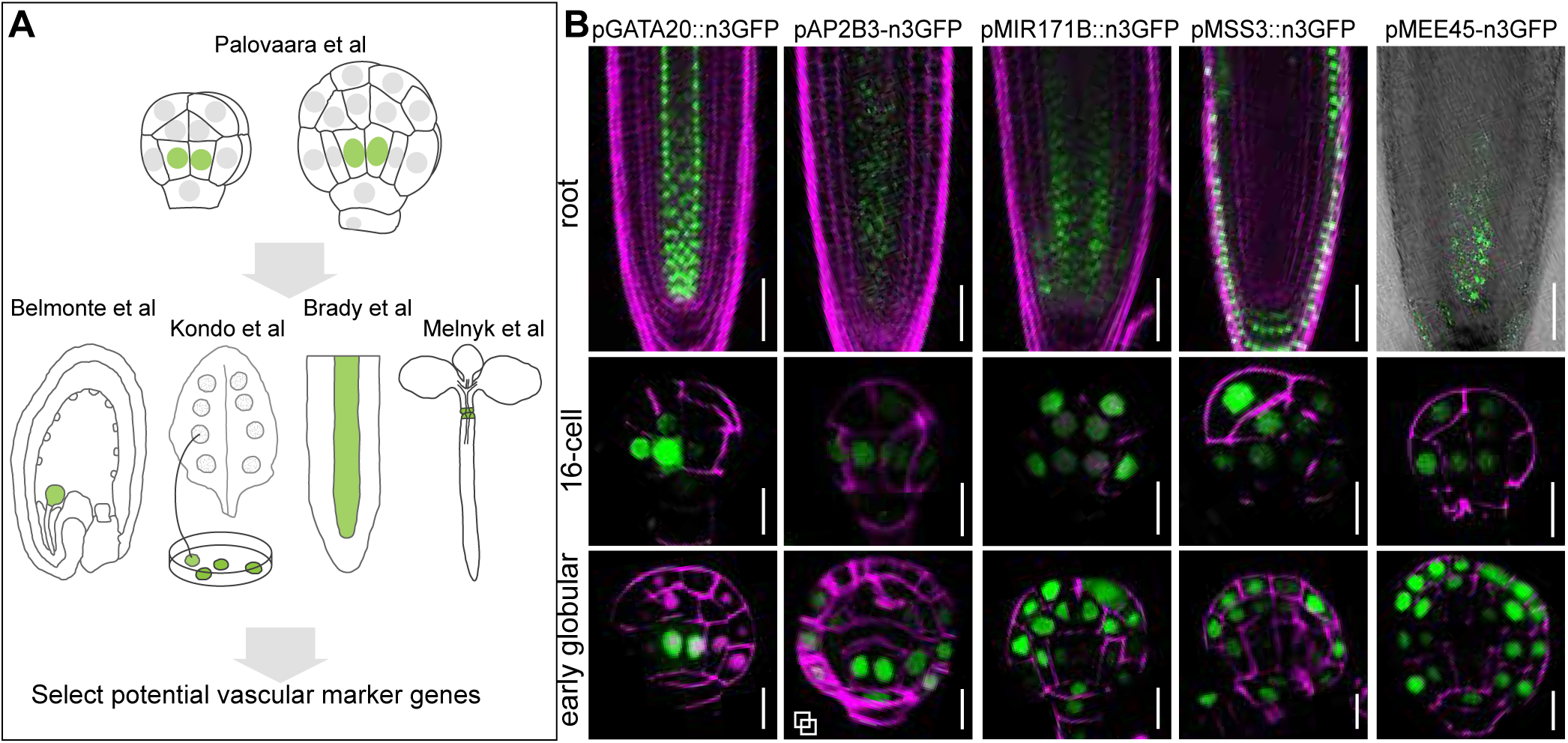
Identification of novel vascular reporters. (A) Overview of transcriptomics datasets used for the selection of new vascular reporters. (B) Expression patterns of new vascular reporters for the embryo. Fluorescent protein signals are displayed in green, cell wall staining in magenta. Roots are stained with PI, embryos with Renaissance. Scale bars represent 50 µm (root) or 10 µm (embryo).

Beyond resolving the ontogeny of vascular tissue initiation in the embryo, our detailed analysis of vascular-specific markers also shows that there are significant differences in gene expression within the vascular tissue between embryo and root. Hence, initiation and maintenance of tissue identity seem to be associated with different gene sets.

### Auxin signaling through MP is necessary, but not sufficient for initiation of vascular identity

Auxin signaling plays many key roles in plant development and one of the clearest is its contribution to vascular development. We sought to investigate the role of auxin signaling in the initiation of vascular identity in the early embryo. To this end we expressed the non-degradable *bdl* protein to block MP activity (Hamann et al. 1999, Weijers et al. 2006a), while examining markers of vascular identity (Figure 3A). Since *bdl* expression in the entire embryo results in early developmental defects (Rademacher et al. 2011, Yoshida et al. 2014), we employed twocomponent gene activation and selectively expressed *bdl* in vascular cells using the Q0990-GAL4; UAS-erGFP driver line (Haseloff, 1999). The GAL4 driver in the Q0990 line is inserted near the ZLL gene (Radoeva et al., 2016) and erGFP thus reports *ZLL* expression. Vascular markers were introduced into the Q0990 background and crossed with a line containing GAL4-dependent UAS-*bdl* (Weijers et al., 2006). The domain of *bdl* expression is marked by ER-localized GFP, whereas the vascular markers are reported by nuclear GFP fluorescence (Figure 3A).

**Figure 3:**
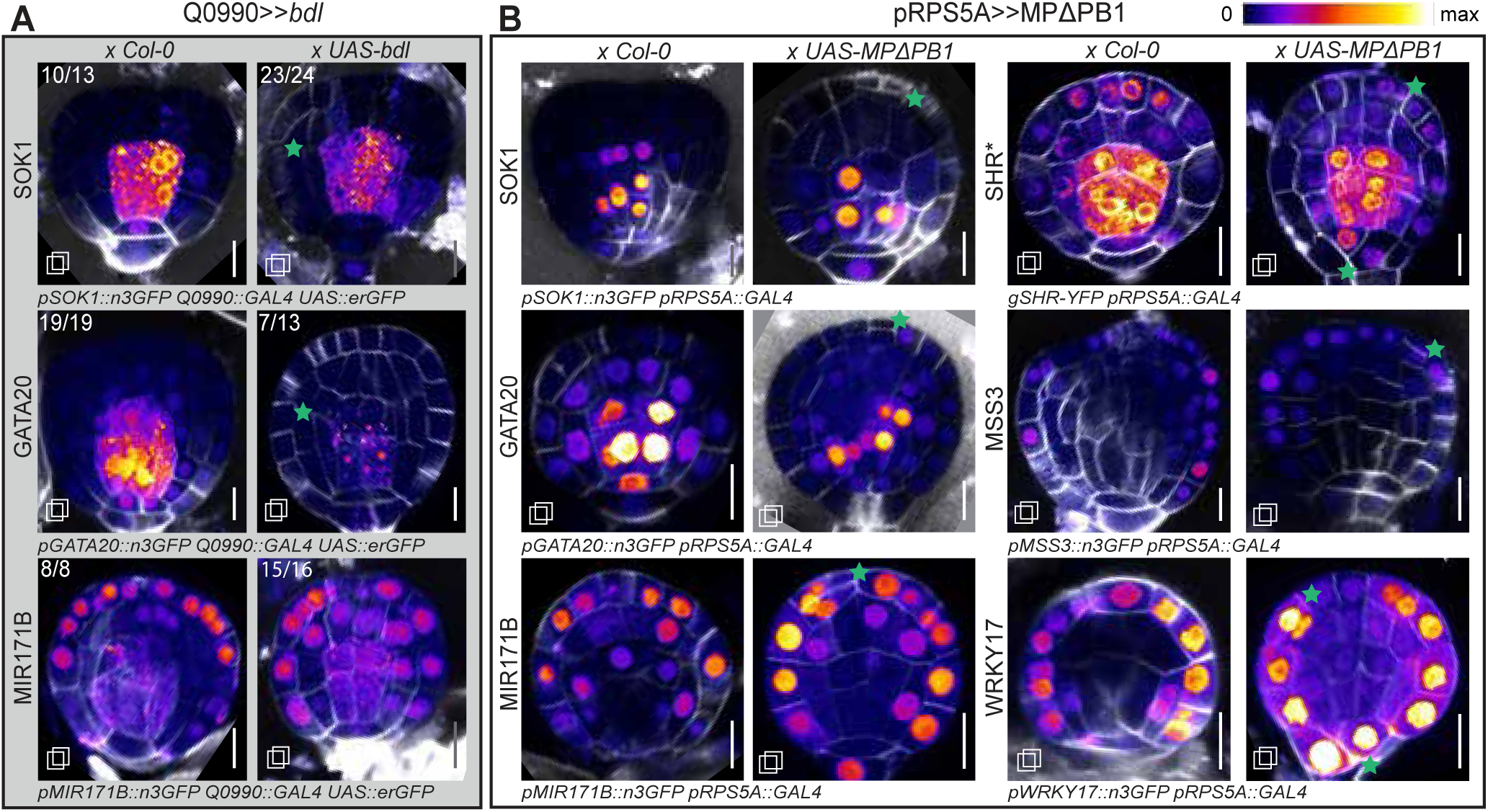
The role of auxin response in embryonic vascular gene expression. (A) Embryos resulting from crosses between a line containing Q0990::GAL4, UAS::erGFP, and a vascular reporter; and either Col-0 or a line containing UAS::*bdl*. Numbers in top left corner of each panel indicate the fraction of embryos observed with the pattern displayed. (B) Embryos resulting from crosses between a line containing pRPS5A::GAL4 and a vascular reporter: and either Col-0 or a line containing UAS::MPΔPB1. Fluorescent protein signals are visualized using the Fire LUT (see inset), embryos are stained with Renaissance (white). Scale bars represent 10 µm. □ Indicates images result from a stack. Note that in (A), vascular reporters are represented by nuclear signal, while the expression domain of the Q0990 reporter (UAS-erGFP) is marked by ER-localized GFP.

Embryos where *bdl* was expressed in the vascular cells often showed altered ground tissue division orientation as previously reported (Möller et al., 2017)(Figure 3A, green asterisks), indicating that auxin signaling was successfully inhibited. However, erGFP expression was not affected in *bdl*-expressing embryos (Figure 3A), indicating that maintenance of *ZLL* expression in presumptive vascular cells does not depend on auxin response. Crosses of the Q0990-GAL4 line with Col-0 wild-type resulted in no such changes in division orientation. *bdl* expression led to 96% (n=24) of the observed embryos lacking nuclear *SOK1* expression whereas wild-type crossed embryos nearly all showed normal *SOK1* expression. As *SOK1* is regulated by MP (Möller et al., 2017; Yoshida et al., 2019), this further confirms the repression of MP activity. However, not all vascular characteristics are gone from the inner cells. In Q0990>>*bdl* embryos, *GATA20* expression is absent in about half (54%, n=13) of the embryos, but remains present in the other half, indicating that in many embryos, repression of vascular identity is incomplete, as is supported by normal *ZLL* expression. In addition, expression of the inverse marker *MIR171B* was mostly unchanged in Q0990>>*bdl* embryos (Figure 3A). These findings indicate that when auxin signaling is blocked in inner cells, vascular identity is compromised, but not abolished. The remaining vascular program is insufficient for further proliferation and development, thus we conclude that auxin signaling through MP is essential for functional vascular tissue specification.

After confirming that auxin signaling is required for vascular initiation, we next asked whether it is also sufficient. While differences in auxin activity across cell layers in the early embryo, as measured by the R2D2 and *DR5v2* reporters (Liao et al., 2015), are small, there is a clearly defined gradient with high levels in central and lower levels in peripheral cells (Möller et al., 2017) (Figure S2). We asked if this small difference in auxin signaling between central and peripheral cells in the embryo is sufficient to restrict (vascular) identity to inner cells. We therefore expressed a version of MP that cannot be inhibited by auxin-dependent Aux/IAA proteins, and that is hyperactive (MPΔPB1)(Krogan et al., 2012) from the ubiquitous *RPS5A* promoter (Weijers et al., 2001) using the GAL4-UAS system. Embryos with ubiquitous MPΔPB1 expression often showed altered division planes in epidermal cells and occasionally in the hypophysis (Figure 3B, green stars), indicating effectiveness of transgene expression. However, ectopic MPΔPB1 expression did not induce ectopic vascular marker expression. Expression of vascular genes (*GATA20*, *SHR*, *SOK1*) remained restricted to the vascular cells and likewise, inverse markers of identity (*MIR171B*, *MSS3*, *WRKY17*) were still expressed exclusively in surrounding cells (Figure 3B). These results show that ectopic auxin response can induce changes in cell division orientation, but is insufficient for inducing vascular tissue specification in the early embryo. This suggests that unknown additional factors limit the domain of vascular identity.

### Identification of transcriptional regulators of vascular gene expression

Given the co-expression of vascular marker genes in the embryo and in the post-embryonic vasculature, it is likely that there are common regulators. To identify transcription factors that bind multiple vascular gene promoters, we performed an enhanced Yeast One Hybrid (eY1H) assay (Gaudinier et al., 2011; Reece-Hoyes et al., 2011). Promoters from 14 of the aforementioned vascular reporter genes were screened against a custom collection of 2037 transcription factors and other DNA-binding proteins in an all-by-all setup (Table S1). Among the 32,592 interactions tested, 1,111 were positive. Combining all interactions resulted in a network comprising 14 promoters and 382 transcription factors (Figure S3, Table 1). This network contained a large number of transcription factors that could bind to many of the vascular promoters. For in-depth analysis, we therefore selected the 6 vascular promoters that each had more than 25 interactors and which showed the most prominent vascular specificity in the early embryo (Figure 4). These 6 consist of: 3 vascular specific marker genes (*GATA20, SOK1, ZLL*); and 3 vascular inverse marker genes (*MIR171B, MSS3, WRKY17*). The network that contains these promoters and their interactors comprises 221 transcription factors and 521 interactions (Figure 4A). If there is a common vascular transcriptional program, it is likely that multiple vascular genes are regulated by a common set of transcription factors. We therefore parsed the interaction network to identify such common transcription factors as potential regulators of vascular identity.

**Figure 4:**
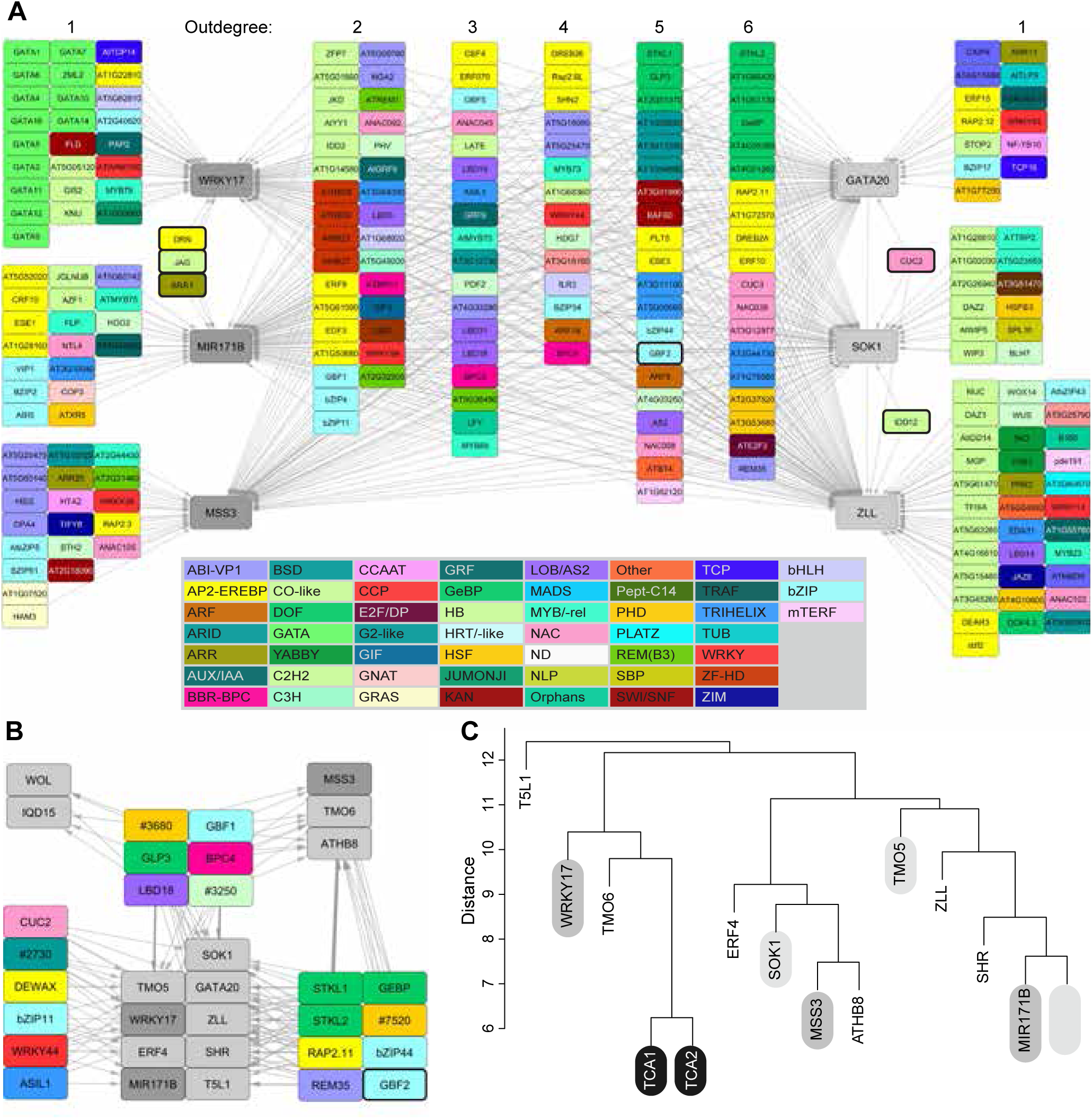
Partial Yeast One Network and selection and candidate regulators of vascular identity. (A) Yeast One Hybrid network showing all interactors of 6 out of 14 vascular promoters screened. Nodes representing transcription factors are colored according to their transcription factor family (see inset) and are grouped by outdegree. Nodes representing promoters are colored light (vascular specific) or dark (vascular inverse) grey. (B) Network overview of the 20 candidate regulators of vascular identity with all 16 vascular promoters screened. Colors as in A. (C) Dendrogram resulting from hierarchical clustering of promoters by interactor set. Branch length indicates distance/similarity in interactor set. Two promoters from an unrelated screen (TCA1/2) were included as an outgroup.

**Table 1:**
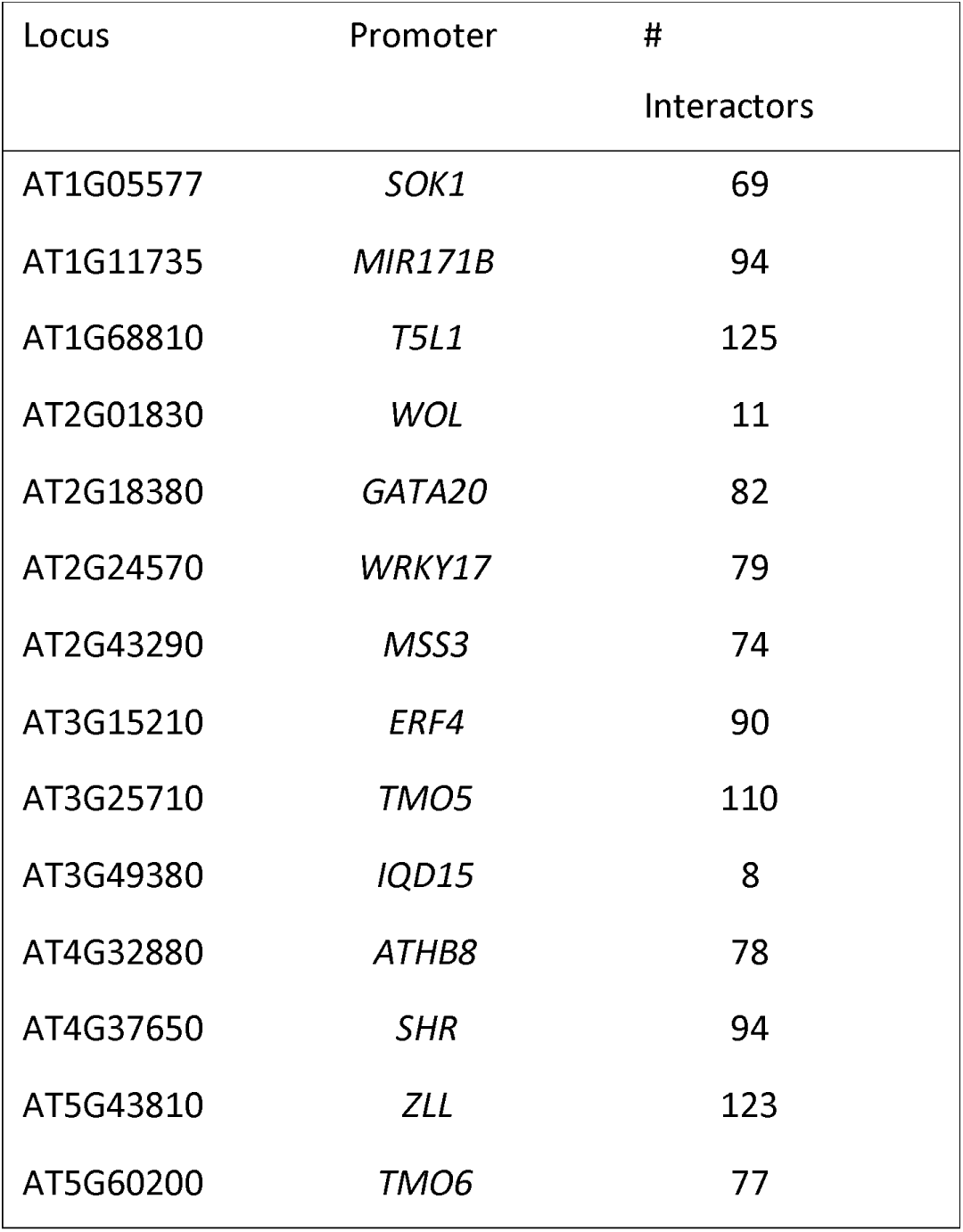
Number of interactions recorded per promoter screened.

A large number of transcription factors were identified to interact with the majority of these promoter sequences. And the majority of these transcription factors bind to both sets of promoters (vascular specific and inverse), only a few can bind to only one set. CUC2 and IDD12 can bind to 3 or more vascular specific markers, but no inverse markers. JAG, DRN and ARR1 bind to 2 or more vascular inverse markers, but no vascular specific markers (Figure 4A). However, these transcription factors are a small minority: in general, the vascular-specific and -inverse promoters have highly similar sets of transcription factors binding to their promoters despite having very different expression patterns. When we perform clustering on the promoters based on their interactor set we find that vascular-specific and vascular-inverse markers do not have distinct sets of interactors (Figure 4D). These findings suggest a large set of transcription factors that could act in complex GRN controlling vascular identity. It should be noted though that both sets of promoters show differential expression between vascular and non-vascular cells, and it is well possible that the same transcription factors (or related proteins) could act as vascular-specific activator or repressor.

To identify candidate regulators of vascular identity, we selected 20 transcription factors from the network. This selection was performed in two steps. First, transcription factors were discarded that (i) bound to few vascular promoters; (ii) were expected to be false positives based on their promiscuous binding profiles in previous screens (Gaudinier et al., 2018; TaylorTeeples et al., 2015); or (iii) were most likely not expressed during embryogenesis based on transcriptomics data (Belmonte et al., 2013; Schlereth et al., 2010). This approach resulted in a list of 50 transcription factors. Next, each transcription factor was scored on: (i) expression during vascular development and early embryogenesis; (ii) number of vascular promoters bound; (iii) diversity of expression patterns bound; and (iv) vascular promoter binding in published DAPseq data (O’Malley et al., 2016). For further analysis we selected the top 20 transcription factors ranked by cumulative score (Table S3).

A candidate regulator can only contribute to vascular identity if that candidate is present at the time and place that identity specification takes place. To ascertain the presence of candidate regulators during embryogenesis, translational fusions of genomic fragments to YFP were created and observed for 17 of these 20 transcription factors (Figure 5A). These revealed that 10 candidate regulators were indeed present at 16-cell stage in the pro-embryo (Figure 5A). The remaining 7 were either not detected during embryogenesis or not at the correct time or location (Figure S4). The majority of the 10 candidate regulators expressed in 16-cell embryos are present uniformly in the nucleus, except for members of the GeBP family, which accumulate in foci within the nucleus, similar to previous reports (Figure 5A)(Curaba et al., 2003). No conspicuous differences between cell types could be found in the early embryo, neither in protein quantity nor localization. This indicates that if these candidates contribute to specifying vascular identity, their cell-specific action is not the result of protein level or location. Instead an unknown mechanism might contribute to cell-specific activity.

**Figure 5:**
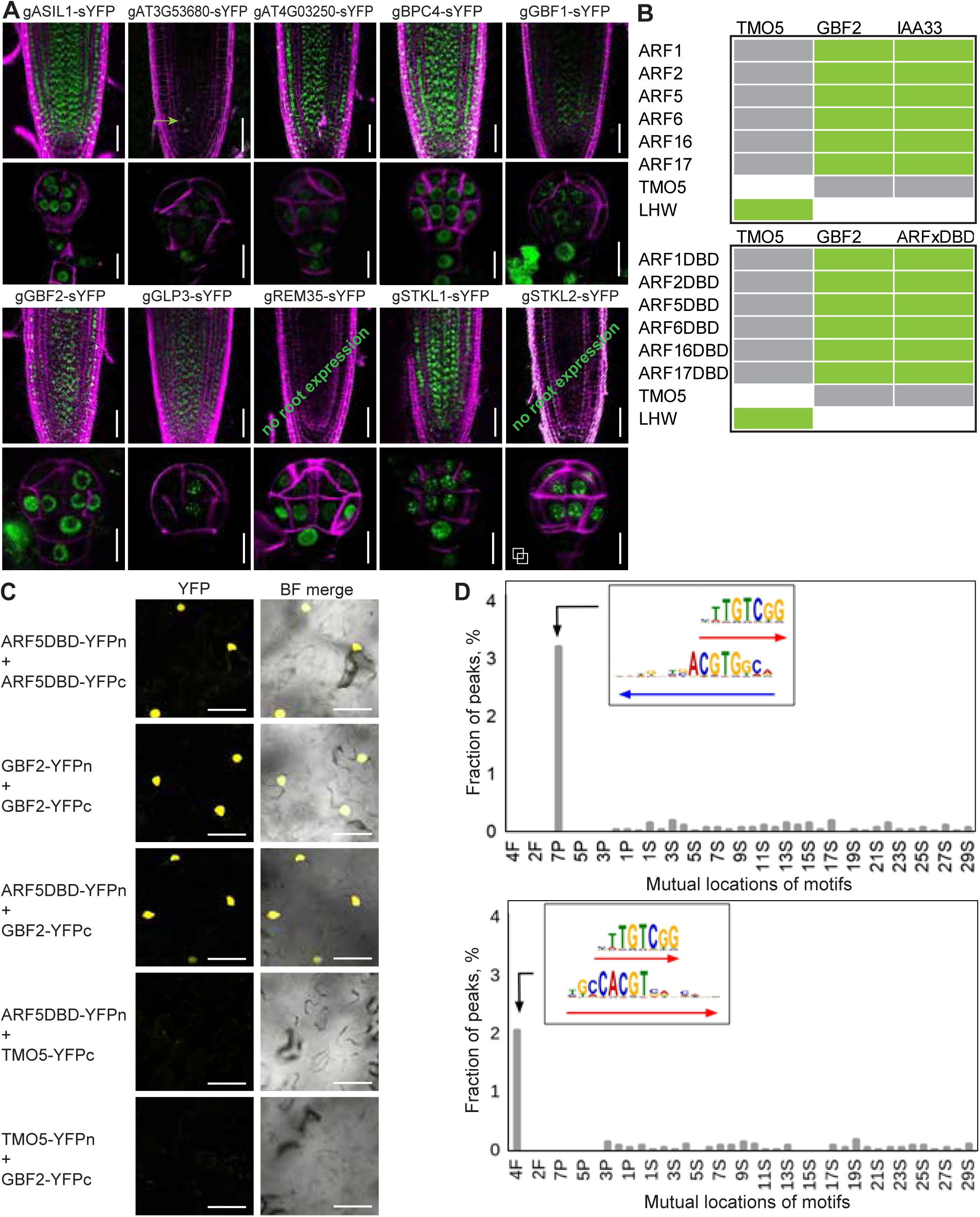
Protein localization and interactions of candidate regulators of vascular identity. (A) Translational reporter lines of 10 candidate regulators in the root tip and early pro-embryo. Fluorescent protein signals are displayed in green, cell wall staining in magenta. Roots are stained with PI, embryos with Renaissance. Scale bars represent 50 µm (root) or 10 µm (embryo). (B) Selected images of split-YFP (BiFC) assays showing the interaction between the DBD of ARF5/MP and GBF2. (C) Overview of split-YFP (BiFC) results indicating that GBF2 can interact with the full-length protein and DBD of 6 different ARFs. (D) Distribution of potential ARF5/bZIP68 composite elements within ARF5 binding regions taken from Dap-Seq. X axis numbers reflect number of nucleotides, F - full overlap, P - partial overlap, S - spacer. Left: ARF5/bZip68 everted composite element distribution. Right: ARF5/bZIP68 direct composite element distribution.

### GBF1 and GBF2 can interact with MP

To determine whether any of the candidate regulators could induce or repress gene activity during vascular specification, each was expressed in meristematic cells with the *RPS5A* promoter (Weijers et al., 2001), either as native cDNA or as a fusion with a dominant SRDX repressor motif. Misexpression of several resulted in either lethal or mild developmental phenotypes, but none of the identified candidate regulators suppressed or ectopically induced vascular tissue formation or differentiation (Figure S5). Therefore it is unlikely that any of these candidates acts in isolation to control vascular tissue initiation. More likely is the control of identity by a complex Gene Regulatory Network (GRN), in which the unique interactions between individual regulators provide cell-type specificity.

In our efforts to understand the mechanisms of gene regulation by MP, we had immunoprecipitated MP-containing protein complexes from root tips in an MP-GFP transgenic line whose functionality had been validated (Schlereth et al, 2010). In this experiment, we identified GBF2 as a potential interactor of MP-GFP (Table S2). As both are candidate regulators of vascular identity, we decided to follow up on this interaction. Immunoprecipitation of GBF2-YFP and its homolog GBF1-YFP, followed by mass spectrometry did not recover MP, presumably due to the very low abundance of MP, but did confirm previous observations that G-class bZIP proteins can heterodimerize extensively (Figure S6). To test MP-GBF interactions more directly, we performed split-YFP assays (BiFC) (Hu et al., 2002; Walter et al., 2004) in *Nicotiana benthamiana*. Both GBF2 and GBF1 could interact with MP (Figure 5B-C, Figure S7). Interestingly, this interaction was not restricted to MP: ARFs of all three major classes (A/B/C: (Okushima, 2005; Finet et al., 2013)) could interact with both GBF2 and GBF1 (Figure 5B, Figure S7), and the interaction domain was mapped to the ARF DNA-binding domain (Figure 5B-C, Figure S7).

GBF proteins were reported to be involved in the responses to blue light and in leaf senescence (Singh et al., 2012; Smykowski et al., 2010; Mallappa et al., 2006; Giri et al., 2017). However, *gbf1*, *gbf2* and *gbf3* single and double mutants show no developmental phenotypes (Figure S7B). This is likely a result of genetic redundancy: the bZIP G-class contains 5 members and double mutants show increased expression of close homologs (Jakoby et al., 2002; Dröge-Laser et al., 2018)(Figure S7B-C). A triple mutant could not be recovered from plants homozygous for *gbf1* and *gbf3*, and segregating *gbf2*, suggesting that lack of all three proteins may result in lethality. Indeed, disruption of a GRN underlying vascular identity establishment would likely result in early developmental arrest. Overexpression using *RPS5A* or *35S* promoters caused pleiotropic developmental defects. pRPS5A>>GBF2-SRDX plants were often sterile, while 35S::GBF1/2 plants had round leaves and showed delayed flowering (Figure S5, S8D). However, no ectopic vascular development was observed. These findings indicate that GBF1/2 protein quantity alone does not specifically limit vascular development.

### GBFs bind to Gboxes close to AuxREs to modulate auxin responsive expression

GBF2 and GBF1 physically with the DNA-binding domain of ARFs, and thus have the potential to co-regulate auxin-responsive genes. Interactions among transcription factors can lead to cooperative DNA binding if both transcription factors can bind to cognate DNA elements in close proximity. Indeed, Gbox motifs were found to be enriched in close proximity to AuxREs (Weiste et al., 2014;Ulmasov et al., 1995) and they are overrepresented in auxin-responsive and ARF-binding regions (Berendzen et al., 2012; Cherenkov et al., 2018). To test the co-occurrence of ARFs and G-class bZIPs motifs we applied the MCOT package (Levitsky et al., 2019) to ARF5 and ARF2 peaks taken from genome-wide DAP-seq profiles (O’Malley et al., 2016). We analyzed all possible combinations of AuxREs (ARF2/5 motifs) and Gboxes (GBF3, bZIP16/68 motifs) with any overlap or spacer lengths below 30 nucleotides, and found that bZIP68 and ARF5 motifs overlap (p-value < 5E-6) (Figure 5D)(Cherenkov et al., 2018).

To investigate the function of these linked motifs, we selected three vascular promoters that contained clear AuxRE and Gbox motifs in close proximity: with an overlap (WRKY17), a short spacer (TMO5) and a long spacer (GATA20) (Figure 6A). Transcriptional reporters with mutated promoters were generated to determine the contribution of the AuxRE-Gbox motif to vascular gene expression domain and level. Removing the complete AuxRE-Gbox motif from the promoters of *GATA20, TMO5* and *WRKY17* resulted in a strong and significant reduction of fluorescence in transgenic roots (Figure 6C-E). For the *WRKY17* reporter, this reduction was more significant in the vascular bundle compared to the rest of the root meristem, resulting in a changed vascular/non-vascular signal ratio (Figure 6E-H). This suggests that for this promoter, the overlapping AuxRE-Gbox motif controls expression levels specifically in vascular cells. Removing only the Gbox had a smaller effect (Fig 6C-E). Mutated *GATA20* and *TMO5* promoters in which the adjoining Gbox had been removed did not show a significant decrease in fluorescence but instead resulted in increased variation in expression level among transgenic lines (Figure 6C-D). Thus the Gboxes in these promoters appear to be contributing to the expression stability instead of absolute vascular expression levels. Since members of several transcription factor families can bind to Gboxes (Schindler et al., 1992; Lian et al., 2017; O’Malley et al., 2016) it is unclear whether GBF2 is the factor contributing to vascular expression stability.

**Figure 6:**
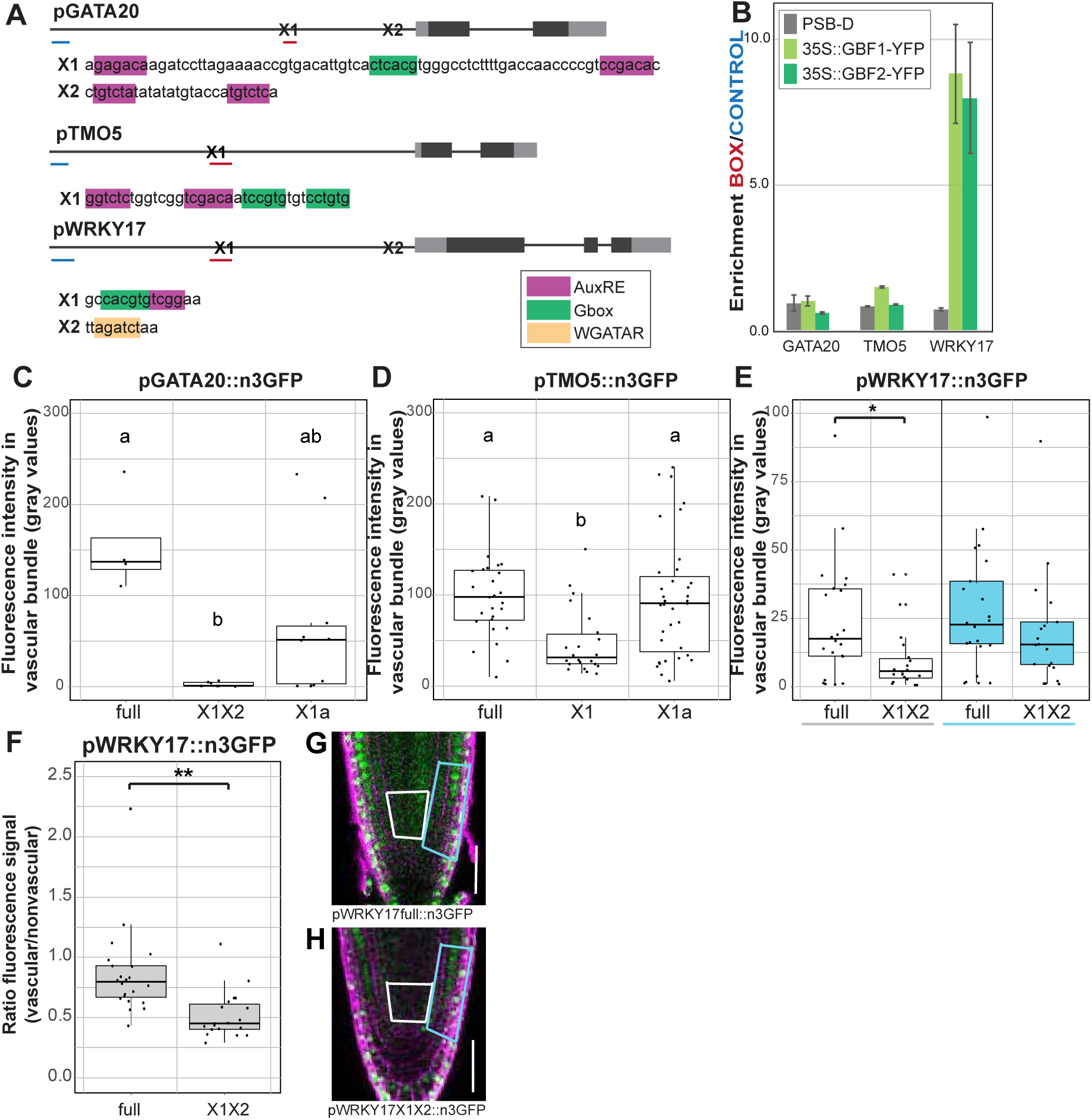
G-boxes can be bound by GBF2 and are needed for stable vascular expression.. (A) Schematic overview of the promoter sequences of *GATA20*, *TMO5* and *WRKY17*. X1 and X2 indicate regions containing TF-binding sites. Blue and red lines indicate control regions and Gbox regions used for ChIP-qPCR. (B) ChIP-qPCR performed on Arabidopsis cell cultures expressing either GBF1-YFP or GBF2-YFP. Relative enrichment of the BOX regions compared to CONTROL regions. Scale bars represent standard error. (C-E) Boxplots displaying fluorescence intensity of transcriptional reporter lines. Each plot compares the mean fluorescence in the measured cells for T1 roots containing full length or truncated promoters of GATA20 (C), TMO5 (D) and WKRY17 (E). Each point is the mean fluorescence in the early vascular cells measured from 1 independent T1 root. For the WRKY17 promoter two areas were measured, the vascular bundle (white) and adjacent non-vascular cells (blue). (F) Ratio of WRKY17 driven GFP signal in the vascular cells compared to signal in the non-vascular cells. (G-H) Expression patterns in representative WRKY17 T1 roots, boxes indicate the region in which fluorescent signal was measured. Scale bars represent 50 μm.

Next, we tested if GBF2 alone can bind to the Gboxes present in vascular promoters. ChIP-qPCR on suspension cell cultures overexpressing GBF2-YFP confirmed that GBF2 can bind to the Gbox motif in the *WRKY17* promoter, but could not confirm the same for the *GATA20* and *TMO5* promoters (Figure 6B). Instead it is likely that that GBF2 and MP both need to be present to interact with the promoters of these two genes.

If GBF2 can bind to vascular promoters and co-regulate vascular gene expression, its overexpression should affect the regulation of vascular genes. To test this directly, we generated protoplasts from vascular reporter lines p*VASC::*n3GFP, and transfected these with a combination of 35S::GBF2-mTurquoise, 35S::MPΔPB1-mScarlet-I and corresponding empty vectors to determine their effects on target promoter activity (Figure 7D-F). MPΔPB1 was used to overcome any auxin-dependent inhibition. Overexpression of only GBF2-mTurqouise had no effect on the promoter activity of *GATA20, TMO5* and *WRKY17*. In contrast, expression of MPΔPB1-mScarlet-I resulted in increased *TMO5* promoter activity and decreased *WRKY17* promoter activity (Figure 7B-C). This effect disappeared when GBF2 was co-expressed with MPΔPB1, suggesting that GBF2 acts by restricting MP activity, probably via competitive binding with the overlapping AuxRE-Gbox motif. This was not the case for the GATA20 promoter, whose activity was increased by both MPΔPB1-mScarlet-I and by MPΔPB1-mScarletI combined with GBF2-mTurqouise (Figure 7A). These findings suggest that the interaction between GBF2 and MP depends on promoter context, yet reflects a functional interaction that contributes to the regulation of vascular genes.

**Figure 7:**
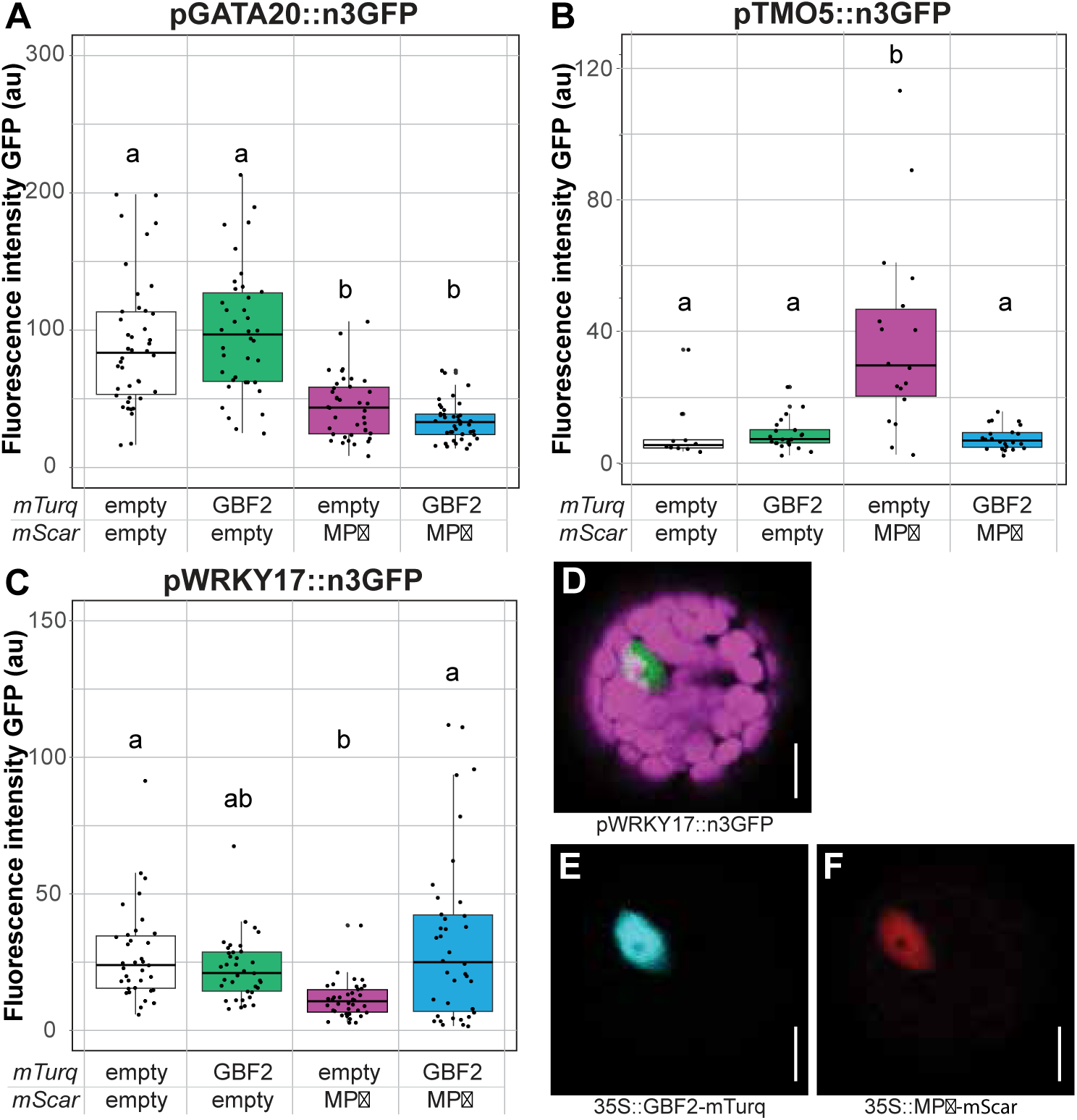
GBF2 modulates MP Δ induced vascular gene expression. (A-C) Boxplots displaying intensity of green fluorescence in nuclei of protoplasts transformed with two misexpression constructs, either empty or containing GBF2 or MPΔPB1. Protoplasts were generated from leaves that expressed pGATA20::n3GFP (A), pTMO5::n3GFP (B), or pWRKY17::n3GFP (C). * indicates p<0.05, ** indicates p<0.001 as calculated by a two-sided Student’s ttest. One-way analysis of variance (ANOVA) followed by Tukey’s honest significant difference (HSD) test was performed to compare either changes in fluorescence intensity, samples were classified into up to 3 categories per experiment (a/ab/b). Results from the Tukey’s HSD test are listed in Table S4. Each assay was performed twice with similar results (data not shown). (D-F) Fluorescence signals detected in protoplast assays. Chloroplasts in pink, GFP in green, mTurquoise in cyan, mScarlet-I in red. Scale bars represent 75 μm.

## Discussion

Vascular tissues play a central role in plant development. Anatomical, physiological and genetic studies contribute to our understanding of vascular development and key aspects of its regulation. While significant insight has been gained into the regulation of cell identity specification, pattern formation, growth and differentiation within the vascular tissue, a key unresolved question is how this tissue is initially specified from non-vascular precursor cells. From lineage tracing in Arabidopsis, it is clear that the vascular lineage has its origins in the early embryo (Scheres et al., 1995), but the origin of embryonic vascular tissue has thus far not been characterized molecularly. Here we have used a panel of vascular marker genes to map the ontogeny of vascular identity from the embryo to post-embryonic tissues. Firstly, we trace its initiation to the 16-cell stage embryo. At this stage the outer cells acquire protoderm identity (Abe et al., 2003) and we find that the inner cells express multiple vascular marker genes, identifying these cells as the first with vascular attributes. These findings are in line with results of a recent transcriptome study which found that the transcriptome of these (inner) cells is similar to that of the later vascular cells (Palovaara et al., 2017). No ground tissue markers have so far been found to be present in the inner lower tier cells (unpublished). This suggests that instead of vascular and ground tissue identities emerging simultaneously, the first ground tissue cells are the daughters of the first vascular cells.

We found that the transcriptional dynamics and progression are vastly different among genes that later mark the vascular domain. This suggests that vascular identity is not a uniform trait that exists across developmental stages, forcing us to reconsider how we view the development of cell type identity over time. Features that distinguish embryonic vs. post-embryonic vascular cells are the co-expression of xylem- and phloem-specific marker genes, and the lack of expression from vascular inverse markers. This suggests that vascular identity initiation as established in the embryo is a temporary state that does not persist. Indeed, in the post-embryonic vascular cells different vascular cell types are highly divergent, a trait which is emphasized by recent advances single-cell (sc)RNAseq. In single-cell RNA sequencing experiments performed on roots, xylem and phloem cells form distinct clusters, but vascular tissues as a whole do not form a cluster that is separated from the two other ‘major’ tissue identities: ground tissue and epidermis (Ryu 2019, Shulse 2019, Denyer 2019). It is questionable if cells in the post-embryonic vascular bundle have or need a common identity. Instead, the brief existence of a “general” primordial, multipotent vascular identity may only be needed when new vascular bundles are initiated: to ensure proper placement of the vascular bundle as a whole and to establish the cambial cells. Future scRNAseq experiments on the embryo, and for example on wounded stems or graft junctions (Melnyk et al., 2018), could help address this question.

There is a strong connection between *de novo* vascular tissue formation and auxin activity. Lack of auxin signaling results in impaired vascular tissue development, while application of exogenous auxin can induce the formation of new vascular bundles (Sachs, 1969; Krogan et al., 2012; Donner et al., 2009; Jacobs, 1952). Auxin and its key transcriptional effector MP are therefore often regarded as master regulators of vascular development (Brackmann et al., 2018). Here we asked if this prominent role also pertains to the earliest steps in vascular tissue specification in the embryo. We find that auxin signaling in the central cells of the embryo is indeed required for the complete establishment of the vascular transcription program. This conclusion is supported by an earlier transcriptome analysis, where many vascular genes were downregulated in embryos where the auxin response inhibitor *bdl* was expressed in the inner cells marked by the Q0990 driver (Radoeva 2016). Interestingly though, several vascular marker genes, including the *ZLL* gene, persist even upon auxin response inhibition. This suggests that for part of the vascular program, auxin response is not required after the initial specification event. Conversely, we find that auxin signaling through MP is not sufficient to induce vascular identity outside of the normal vascular domain. It should however be noted that a dominantactive version of MP lacking its C-terminal PB1 domains was used. It is possible that interactions with the PB1 other than Aux/IAA inhibition – such as for example homooligomerization (Nanao et al. 2014) – are required for MP’s activity in vascular development. Nonetheless, this result suggests that in addition to auxin response, there must be additional, yet undiscovered factors that determine which cells acquire vascular identity. Given the small size of the embryo, a system built on a set of regulators would also provide a more robust mechanism for the regulation of identity than a single master regulator of identity.

Using an enhanced Yeast One Hybrid (eY1H) assay we identify 10 transcription factors that can bind to a large number of vascular promoter sequences and which are also present at the moment of vascular identity specification. While the exact roles and relations of these transcription factors remain unclear, they could be part of the GRN controlling vascular initiation and provide a new entry point into studying vascular tissue specification. A complex GRN would provide a robust system for the initiation of identity. And while it is clear that auxin signaling through MP is necessary, other candidate regulators of identity have not yet been untangled. As single perturbations of other components had little effect we decided to explore the link between MP and a new candidate regulator: GBF2 and its close homolog GBF1.

GBF1 and GBF2 not only bind multiple vascular gene promoters, but they also interact with the DNA-binding domains (DBDs) of ARF proteins of all three major classes (A,B,C; (Okushima, 2005; Finet et al., 2013)). Other previously identified ARF-interacting proteins were shown to interact with the PB1 domain (Ripoll et al. 2015, Shin et al. 2007, Varaud et al. 2011) or middle region (Wu et al., 2015), which would likely modulate transcription activity. Instead, interaction of GBFs with the DBD could modify DNA-binding properties, by exclusion or cooperativity. GBF binding motifs, Gboxes, were often found in close proximity to AuxREs (Berendzen et al., 2012; Cherenkov et al., 2018; Ulmasov et al., 1995), suggesting that GBFs and ARFs could regulate gene expression together. We show that the Gboxes in several vascular promoters affect the stability of expression levels in the vascular bundles and that GBF2 was able to prevent the effect of MPΔPB1 on several target genes. Thus GBF1/2 was able to modulate and/or stabilize auxin-dependent regulation of vascular gene expression.

GBF1/2 is a strong candidate for being part of the GRN that controls the initiation of vascular identity. A *gfb1 gbf2 gbf3* triple mutant could not be recovered, and its lethality highlights a common theme in investigating the regulation of basic cell identities. Further genetic analysis, for example using conditional mutant alleles, could help to define the role of these transcription factors in vascular tissue initiation. The broad expression of GBFs could suggest that, similar to MP (Möller et al., 2017), their predicted cell-specific activity is influenced by local signals, such as redox potential and phosphorylation (Shaikhali et al., 2012; Klimczak, 1992; Smykowski et al., 2016). In a larger GRN controlling vascular identity, GBF1/2 could contribute to limiting vascular identity to the innermost cells of the early embryo, future research into these candidate regulators is needed to confirm such a role for GBF1/2.

In conclusion, our work identifies embryonic vascular tissue identity as a primordial state from which vascular cell types evolve in diverse patterns. We find that auxin response is necessary, but not sufficient for specifying this initial vascular identity. We identify a range of potential regulators of vascular identity and suggest a complex GRN being in control of vascular identity. One potential regulator, GBF1/2, can interact with MP to modulate vascular gene expression. We expect that further analysis of these and other candidate regulators will help identify the elusive mechanism that directs vascular tissue identity.

## Material and methods

### Plant material and growth conditions

All Arabidopsis plants used in this study were of the Col-0 ecotype, except for the cell cultures, which were L*er*. Reporter lines for *DOF6*, *PEAR1* and *TMO6* were previously published in Miyashima et al. (2019). Transcriptional reporters for targets of MP: *IQD15*, *SOK1*, *T5L1*, *TMO5* and *WRKY17* were previously published in De Rybel et al. (2013), Möller et al. (2017) and Schlereth et al. (2010). The reporters for *ATHB8* and *SHR* were previously published (Donner et al. 2009, Nakajima et al. 2001). New reporters were generated for *WOL* and *ZLL* using primers documented in Table S5, and these reproduce previously described expression patterns (Mähönen et al., 2000; Radoeva et al., 2016). All newly generated transcriptional and translational reporter constructs (see below) were transformed into the Arabidopsis Col-0 accession. Misexpression lines were generated by introducing *UAS-gene* contructs into a background containing the *pRPS5A-GAL4* driver or by introducing *35S*-driven constructs into the Col-0 background. T-DNA insertion lines *gbf1* (SALK_027691), *gbf2-1* (SALK_206654), *gbf2-2* (SALK_205706), *gbf3* (SALK_067963) were obtained from the Arabidopsis stock centers (NASC and ABRC). Plants were genotyped using the primers listed in Table S5. Arabidopsis seeds were surface-sterilized, plated on half-strength Murashige and Skoog (MS) medium with the appropriate antibiotic (50 mg/l kanamycin or 15 mg/l phosphinothricin) and underwent 2 days of stratification at 4 °C before being placed in the growth chamber. Plants were grown at 22 °C under standard long-day (16 h light 110 µE m−2 s−1 [Philips Master TL-D HF 50W/840] and 8 h dark) conditions.

### Plant growth methods

Wild type Arabidopsis Landsberg *erecta* and transgenic PSB-D cell suspension cultures were maintained in MSMO medium in the dark at 25°C gently shaking at 130rpm. Cells were sub cultured every 7 days in a 1:10 dilution with fresh medium. Transformations were conducted without callus selection as described by (Van Leene et al., 2007).

Bimolecular Fluorescence Complementation (BiFC) was performed by infiltrating *Nicotiana benthamiana* leaves with *Agrobacterium tumefaciens* strains carrying the appropriate plasmids (see below). Two days after infiltration, leaf sections were cut and imaged by confocal microscopy.

Protoplasts were harvested with a tape sandwich (Wu et al., 2009) and transfection was performed as described in (Russinova et al., 2004). Protoplasts were prepared from plants containing a stable vascular transcriptional reporter. Green fluorescence levels were measured in protoplasts with both red (mScarlet-I) and blue (mTurquoise) fluorescence two days after transfection using confocal microscopy.

### Vector construction for plant transformation

All constructs for plant transformation were cloned using SliCE cloning into previously published LIC vectors (Wendrich et al., 2015; Zhang et al., 2014). All primers used for cloning can be found in Table S5. Promoters for transcriptional reporters were introduced into the pPLV04_v2 backbone (De Rybel et al., 2011). Translational fusion constructs were generated by amplifying up to 3 kb of the promoter and the gene up to but not including the stop codon and introducing this sequence into pPLV16_v2. *UAS-gene-SRDX* overexpression constructs were cloned by introducing the amplified cDNA sequence without stop codon into a modified pPLV32_v2 backbone containing a SRDX peptide using SLICE cloning (Wendrich et al., 2015; Zhang et al., 2014). *35S* overexpression constructs were generated by introducing the cDNA sequence into a modified pPLV26 containing a C-terminal YFP. All constructs were introduced using the simplified floral dip method as described in De Rybel et al., (2011). BiFC constructs were generated by introducing amplified cDNA sequences into modified pPLV26 vectors containing NtYFP or CtYFP either before or after the insertion site. Binary vectors for misexpression in protoplasts were generated by introducing the cDNA of GBF2 or MPΔPB1 (first 766 amino acids) into pMON99 containing C-terminal mTurquoise or mScarlet-I. All primers used for cloning are listed in Table S5.

### Microscopy

Confocal imaging was performed on a Leica SP5 II system equipped with Hybrid Detectors. Confocal microscopy was performed as described previously (Llavata-Peris et al., 2013). Cell walls were visualized by staining with propidium iodide (PI, for roots) or SCRI Renaissance Stain 2200 (Renaissance Chemicals R2200, for embryos) respectively.

### Yeast One Hybrid

Enhanced Yeast One Hybrid assays were performed as described (Gaudinier et al., 2017). The promoter used for the yeast reporter constructs (pMW2 and pMW3) was the same as the promoter used for reporting localization in Arabidopsis. The prey collection used was the complete Arabidopsis transcription factor collection available at the Brady lab in July 2016 (Table S1). Network analysis was performed in Cytoscape (Shannon et al., 2003).

### Affinity purification mass spectometry sample preparation

For affinity purification, either 4 g root material or 50 ml of 3 day old transgenic PSB-D cell suspension cultures was used, and protein extraction, pull-down and sample preparation was performed as described (Wendrich et al., 2017).c Peptides were applied to online nano LCMS/MS (Thermo Scientific) using a 60 minute acetonitrile gradient from 5-50%. Spectra were recorded on a LTQ-XL mass spectrometer (Thermo Scientific) and analysed according to (Wendrich et al., 2017). Maxquant output Proteingroups.txt was filtered in Perseus v1.6.2.3.. Volcano plots were generated in R and further visualized in Adobe Illustrator.

### Motif analysis

Analysis of potential binding sites presence was performed with position weight matrices taken from Plant TFDB database (Jin et al., 2017) for GBF3 (MP00318), bZip16 (MP00291) and bZip68 (MP00173). Colocalization of binding sites with ARF binding sites were analyzed with the MCOT package (Levitsky et al., 2019) using data on ARF2 (GSM1925138, GSM1925826) and ARF5 (GSM1925827) binding regions from Dap-Seq analysis (O’Malley et al., 2016).

### ChIP-qPCR

ChIP-qPCR was performed on Arabidopsis cell cultures using a protocol adapted from (Gendrel et al., 2005). 3-4 grams of filtered cell culture material was used as input material. After crosslinking and DNA fragmentation, the sample was split and GFP-Trap beads (Chromotek) were used to pull down GBF-YFP complexes while Myc-Trap beads (Chromotek) were used for the negative control sample. qRT-PCR was performed using primers listed in Table S5. Ct values were then used to calculate fold enrichment and relative fold enrichment compared to the control regions.

## Ackowledgements

The authors are grateful to Branimir Velinov, Koyan Bruggeling, Surabissree Merialu Diwakar, Sebastien Paque, Frederique Polder, Allison Gaudinier, Anne-Maarit Bagman, Joel Rodriguez-Medina, Pawel Roszak, Michelle Tang and Peter Etchells for help with experiments, Weijers lab members for helpful discussion and to Camila Lopez-Anido for comments on the manuscript. This work was supported by grants from the Netherlands Organization for Scientific Research (NWO; 831.14.003 to M.E.S. and VICI 865.14.001 to D.W.), the European Molecular Biology Organization (EMBO ASTF-368-2016 to M.E.S.), the Russian Foundation for Basic Research (19-44-543006 to D.N.), a Wageningen Graduate Schools Sandwich PhD Scholarship (to D.N.) and a Budget Project (0259-2019-0008 to D.N.). S.M.B. is an HHMI Faculty Scholar.

---

*Figure S1: Diverging expression patterns between root and embryo.* (A) Activity of transcriptional reporters in the root tip and embryo. Fluorescent protein signals are displayed in green, cell wall staining in magenta. Roots are stained with PI, embryos with Renaissance. (B) Expression of S32 in phloem cells and vascular stem cells in the root tip. Scale bars represent 50 µm (root) or 10 µm (embryo).

*Figure S2: Auxin accumulation and signaling output in the early embryo.* 16-cell (A,C) and early globular (B,D) stage embryos reporting the relative amount of auxin or auxin signaling per cell. (A,B) Relative amount of auxin signaling per cell output as reported by pDR5-n3GFP (left) or pDR5v2-ntdTomato (right). (C,D) Relative accumulation of auxin per cell as reported by R2D2. Left: overlay of signals from undegradable pRPS5A-mDII-tdTomato (red), degradable pRPS5A-DII-3xVenus (green) and Renaissance (white). Right: difference between DII signal and mDII signal per pixel. All images are stacks and all scale bars represent 10 μm. □ Indicates images result from a stack.

*Figure S3: Vascular Yeast One Hybrid network.* Network containing all interactors of the 16 vascular promoters screened. Nodes corresponding to promoters are larger and colored grey (dark grey for inverse markers), nodes corresponding to transcription factors (TFs) are placed together based on their outdegree. TF nodes with an outdegree of 1 are placed on the periphery near their target, TF nodes with an outdegree of 2 or higher are placed in the center and are grouped based on their outdegree, nodes with a higher outdegree are located further to the right. Network overview with TF nodes colored according to TF family. Each TF family is represented by a color and number (see insert table).

*Figure S4: Protein localization of the candidate regulators in root and embryo.* In each panel, YFP fusion protein localization in the root is shown at the top, followed by localization in dermatogen stage, early globular stage and one later stage.Scale bar m in roots or 10 μm in embryos.

*Figure S5: Misexpression phenotypes of SRDX-tagged candidate regulators.* Adult plants expressing SRDX-tagged candidate regulators under the RPS5A promoter show altered leaf morphology or decreased fertility.

*Figure S6: IP-MS/MS experiments confirm that GBF1 and GBF2 can heterodimerize with G-class bZIPs.* Results of immunoprecipitation followed by tandem MS (IP-MS/MS) on Arabidopsis cell cultures expressing GBF1-YFP (left) or GBF2-YFP (right) under the 35S promoter compared to wildtype (PSB-D) cell cultures. Volcano plots show fold change (FC, x-axis) and significance (FDR, y-axis) of each detected protein. Proteins with a p-value below 0.05 (-log(FDR)>1.301) and a fold change above 5 are marked and have their name displayed.

*Figure S7: Split-YFP experiments performed using tobacco leaves to confirm GBF-ARF interactions.* (A) Interactions GBF1/2-CtYFP and ARF-NtYFP. TMO5 was used as a negative control, homodimerization was used as a positive control for GBF and IAA33 was used as a positive control for ARFs. (B) Interactions GBF1/2-CtYFP and ARFxDBD-NtYFP. TMO5 was used as a negative control and homodimerization was used as a positive control for GBF and for ARFs. LHW was used as a positive control for TMO5.

*Figure S8: Genetic analysis of GBF function* (A) Insert locations of the T-DNA lines used. (B) 24-day old plants of 3 sets of GBF double mutants, single mutants and background plants. (C) Relative expression levels of GBF1, GBF2 and GBF3 in the gbf1gbf3 double mutant and single mutants. (D) 35S-GBF1/2 and Col-0 plants 38 DAG. GBF1/2 overexpression lines have early leaves with an increased width/length ratio, late leaves with increased serration and more pronounced veins, and slower development resulting in delayed flowering. No obvious changes in venation pattern were observed.

*Table S1: List of the full TF collection used for yeast 1-hybrid screening*.

*Table S2: IP-MS/MS performed on MP-GFP indicates MP interacts with GBF2*.

*Table S3: Ranking of and scores awarded to the 50 initially selected transcription factors.* Left to right: (grey) Final results of the 4 scoring totals and average total ranking. (yellow/pink) Scores awarded based on binding pattern. Includes data on outdegree and binding pattern from the network and DAPseq data from O’Mally. (grey/blue) Scores awarded based on embryo expression. Includes expression percentile data on embryo expression levels of whole WT embryos (Weijers lab) and fold changes from the embryo expression atlas (Palovaara). (green) Scores awarded based on vascular expression. Includes expression fold changes from leaf disk (Kondo) and expression percentile data from root transcriptome atlas (Brady)(Phloem, Stele, Xylem). * HTA2 and ESE3 were excluded.

*Table S4: P-values resulting from Tukey’s tests performed on promoter deletion and protoplast assays*.

*Table S5: Primers used in this study*.

